# Exhausted mature dendritic cells exhibit a slower and less persistent random motility but retain chemotaxis against CCL19

**DOI:** 10.1101/2021.10.11.463881

**Authors:** Yongjun Choi, Vijaya Sunkara, Yeojin Lee, Yoon-Kyoung Cho

## Abstract

Dendritic cells (DCs), which are immune sentinels in the peripheral tissues, play a number of roles, including patrolling for pathogens, internalising antigens, transporting antigens to the lymph nodes (LNs), interacting with T cells, and secreting cytokines. The well-coordinated migration of DCs under various immunological or inflammatory conditions is therefore essential to ensure an effective immune response. Upon maturation, DCs migrate faster and more persistently than immature DCs (iDCs), which is believed to facilitate CCR7-dependent chemotaxis. It has been reported that lipopolysaccharide-activated DCs produce IL-12 only transiently, and become resistant to further stimulation through exhaustion. However, little is known about the influence of DC exhaustion on cellular motility. Here, we studied the cellular migration of exhausted DCs in tissue-mimicked confined environments. We found that the speed of exhausted matured DCs (xmDCs) decreased significantly compared to active matured DCs (amDCs) and iDCs. In contrast, the speed fluctuation increased compared to that of amDCs and was similar to that of iDCs. In addition, the diffusivity of the xmDCs was significantly lower than that of the amDCs, which implies that DC exhaustion reduces the space exploration ability. Interestingly, CCR7-dependent chemotaxis against CCL19 in xmDCs was not considerably different from that observed in amDCs. Taken together, we report a unique intrinsic cell migration behavior of xmDCs, which exhibit a slower, less persistent, and less diffusive random motility, which results in the DCs remaining at the site of infection, although a well-preserved CCR7-dependent chemotactic motility is maintained.

## Introduction

Recent advances in microfabrication have provided an opportunity to observe single-cell level motility in a tissue-mimicking microenvironment. This approach has become an important milestone in learning the correlation between cellular motility and function in many biological phenomena, including cancer metastasis, wound healing, and immune responses, revealing that the surrounding physicochemical environment significantly impacts cell migration.^1, 2^

Dendritic cells (DCs) are professional antigen-presenting cells that play key roles in the immune response function, wherein the precise control of their migration is essential for the maintenance of tissue homeostasis and immune surveillance. Immature DCs (iDCs) patrol the confined environment of peripheral tissues to search for antigens to present to T cells. An intermittent mode of migration showing an alternative switch between persistent and diffusive motility has been observed for iDCs, which may explain how iDCs efficiently search for rare target pathogens in a tissue-microenvironment.^3-6^ As iDCs are activated upon stimulation, matured DCs (mDCs) upregulate major histocompatibility complex (MHC) II molecules and co-stimulatory molecules.

The mDCs also upregulate the chemokine receptor (CCR7), which allows them to respond to the gradients of CCR7 ligands (CCL19 and CCL21) for their directional migration towards lymphatic organs.^7-12^ Although the chemotactic migration of mDCs is relatively well studied, there is still a lack of understanding regarding DC maturation and the motility of mDCs because DC maturation is a complex phenomenon influenced by dynamic physicochemical cues from the external environment.^13, 14^

Upon short-term (e.g., 30 min) activation with lipopolysaccharide (LPS), mDCs are known to exhibit fast and persistent migration compared to iDCs in the absence of a chemokine gradient, which may facilitate the movement of mDCs over more considerable distances to ensure a more efficient space exploration.^15^ Although short LPS stimulation is sufficient for DC maturation in terms of the upregulation of MHC, CCR7, and co-stimulatory molecules, the resulting DCs do not secret cytokines such as IL-12.^13^ In contrast, DCs can become ‘exhausted’ during longer periods of stimulation (e.g., 24 hours), meaning that they stop producing cytokines and become resistant to further stimulation.^16-19^. More specifically, it has been reported that IL-10 and IL-12 are secreted during stimulation, but that this secretion capacity drops after 24 h. In addition, a recent study showed that exhausted CD8 T cells present distinct migration patterns for maintaining their quiescence and stem-like program under the conditions of a chronic viral infection.^20^ However, it is not yet clear whether DC exhaustion of the cytokine secretion capability affects the cellular locomotion of mDCs.

Thus, we herein report our quantification of both the intrinsic and chemotactic motilities of mDCs activated over different LPS stimulation times. To provide a more reproducible confined microenvironment to monitor two-dimensional (2D) motility of DCs, we develop a modified agarose assay chip using a gel confiner. In addition, we employ a microfluidic channel array containing a chemokine gradient to mimic DC migration toward the lymph nodes (LNs).

## Materials and Methods

### Preparation of the bone-marrow-derived DCs

The bone-marrow-derived DCs (BMDCs) were obtained as described in the literature, ^21, 22^ and further details regarding the experimental methodology are provided in the Electronic Supplementary Information (ESI). In brief, murine bone marrow-derived iDCs were used to differentiate between active matured DCs (amDCs) and exhausted DCs (xmDC). To prepare the xmDCs, iDCs were incubated with LPS (100 ng/mL) for 24 h. ^17^ AmDCs were stimulated according to the same method but over an incubation time of only 30 min, followed by washing and incubation for 24 h in fresh complete medium.^15^ All animal experiments were performed under protocols approved by the Institutional Animal Care and Use Committee of Ulsan National Institute of Science and Technology (UNISTIACUC-19-15). Flow cytometry was performed to characterise the DC phenotype (Cytoflex, Beckman Coulter) (**Figs. S1A** and **S1B**). The expression levels of the co-stimulatory molecules (i.e., CD86, CD80, CD40), the antigen-presenting molecule (I-A/I-E), and the chemokine receptor (CCR7) were verified as DC maturation markers, and CD11c and CD11b were tested as DC markers.^22^ Details of the flow cytometry analyses and cytokine measurements are given in the ESI.

### Preparation of the gel confiner for the under-agarose assay

The gel confiner was composed of a custom-designed polydimethylsiloxane (PDMS) structure and low melting agarose gel. Its preparation procedure is provided in the ESI. Approximately 800 cells were seeded on coverslips (10 mm diameter) coated with bovine fibronectin serum (20 μg/mL, FN, Sigma; cell density = 1.0 × 10^−4^ cells/μm^2^). The cells were incubated for 20 min in a cell culture incubator. The cell suspension was carefully washed three times with fresh complete medium and assembled in the gel confiner.

### Live cell imaging

Live cell imaging of the DCs was performed using an inverted microscope (Eclipse Ti-E, Nikon) configured with a 10× dry objective lens and an sCMOS camera (Flash 4.0, Hamamatsu). The cell trajectories were imaged after 6 h of confinement, and bright-field images were obtained every 1 min for 12 h. Live-cell imaging experiments were performed using a custom incubator system (Chamlide HK, Live Cell Instrument) maintained at 37 °C and containing 5% CO2.

### DC motility analysis

Motility analysis was performed using Imaris (Bitplane) or a homemade MATLAB (MathWorks) code modified from the literature.^23^ The instantaneous speed was obtained by dividing the length between two consecutive frames by the corresponding time difference. The mean track speed is the track length divided by the track duration, where the track length is the total length of displacements within the track. The speed fluctuation is the standard deviation of the instantaneous track speed divided by the mean track speed. The track straightness is the track displacement divided by the track length, where the track displacement is the distance between the first and last positions. The travel range is the maximal distance from the track origin to the farthest position of the track.

The cell trajectory was analysed in terms of the mean-square displacement (MSD) and fitted with a power law according to MSD (τ) = 4Dτ^α^, where D is the diffusion coefficient, τ is the time lag, and α is the scaling exponent. The normal Brownian diffusion is indicated by α = 1. Otherwise, the motility is referred to as subdiffusive (α < 1) or superdiffusive (α > 1).

The turning angle distribution was defined as the angle between two consecutive displacement vectors. This distribution provides statistical information regarding the directional movement of cells. If the cells move persistently, the turning angle distribution has a peak at around 0°. If cell motility is random, a broad and uniform distribution of the turning angle is expected in any direction (0°–180°). If the cell motility is spatially anisotropic, there is a high chance of angular displacements along the 0° and 180° directions. In this study, the turning angle distribution was presented in terms of the time lag, which ranged from 1 to 12 min.

### Chemotaxis against CCL19 gradient

The microchannels were prepared as described previously. ^21^ More specifically, microchannels were prepared with dimensions of 20 µm (width) × 3 µm (height). Specific details regarding preparation of the chemotaxis chip are provided in the ESI. To validate the gradient stability, we examined the diffusion of a fluorescence conjugated dextran (CF488A-dextran 10 kDa, Biotium) over time using fluorescence confocal microscopy (LSM980, Zeiss). After cell loading, the images were recorded between 5 to 15 h. Tracks showing displacement that did not exceed 250 μm were excluded from the analysis. The cell trajectories were manually tracked from the entrance to the end of the microchannel.

### Statistical analysis

Statistical analysis was conducted using OriginPro 2020 (Origin). For the normality test, the D’Agostino Pearson omnibus K square test was used for the single-cell scale pooled data, and the Kolmogorov–Smirnov test was used for representative fraction values obtained from independent experiments. In box plots, the bars include 95% of the data, the boxes contain 75% of the data, the centre bars correspond to the median, and the square points represent the mean values. When the data were normally distributed, statistical analysis was performed using a one-way ANOVA, followed by either Tukey’s multiple comparison test or the two-tailed unpaired t-test for post-hoc analysis. When the data were non-normally distributed, the Kruskal-Wallis with Dunn’s test or the two-tailed Mann-Whitney test was used for post-hoc analysis.

## Results

### Use of the gel confiner to provide reproducible DC confinement

The precise control of cell confinement is critical to reproducing the DC motility observed in tissues.^24^ However, the previously reported under-agarose cell migration approach has technical limitations in terms of studying 2D random motility because cells rarely infiltrate under agarose without a directional cue.^25^ Thus, only those infiltrated cells, not a total population, could be included in the analysis resulting a rather biased interpretation specific for a small fration of the cell population. Covering the cells with a bulk gel block could be attempted, but the degree of confinement was not reproducible.^26^ Microfabricated roof structure with pillers could be used to cover the cells, in which the degree of confinement is controlled by the height of the pillars.^27, 28^ However, the cellular locomotion can be hindered by the presence of the pillar arrays and cellular metabolism could be disturbed due to the narrow space between PDMS roof and glass bottom.

Thus, we developed a modified under-agarose assay chip using a gel confiner, which covered all DCs on the surface and allowed easy handling (**Fig. 1A**). The agarose gel present in the confiner provides a tissue-like confinement environment, and the thin sticky layer of PDMS under the rim contributes to stable and precise confinement over a broad area (∼ 78.5 mm^2^) (**Fig. 1B, C**). The cell height was measured using 3D confocal imaging (**Fig. 1D**), and upon comparison between the ‘Not confined’ (7.6 ± 0.3 μm, mean ± S.E.) and ‘Only gel’ (7.9 ± 0.3 μm) systems, it was apparent that the gel confiner enabled a reproducible and long lasting confinement with cell heights of 2.9 ± 0.4 and 3.2 ± 0.4 μm after 1 and 24 h, respectively (**Fig. 1E**). These results demonstrate that the gel confiner can successfully maintain the cells in a broad area of confined space for a sufficient time to observe free movement of the cells over millimetre-scale fields of view (1300 × 1300 μm^2^).

**Figure 1.**
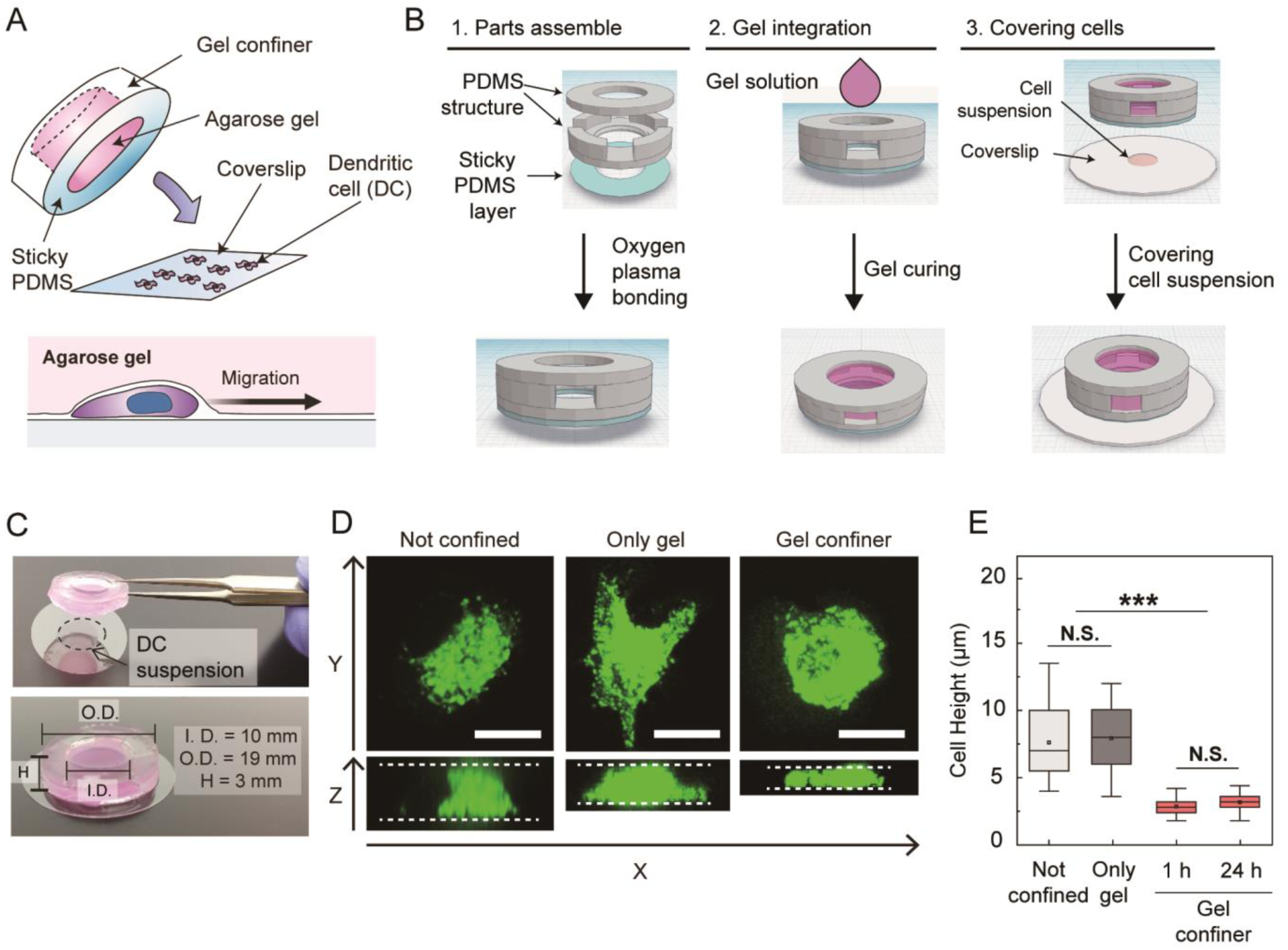
A PDMS gel confiner chip for a more controlled confinement. **(A)** Schematic representation of the modified under-agarose assay based on the gel confiner. **(B)** Fabrication of the gel confiner. **(C)** Photographic image of the gel confiner chip with its corresponding dimensions. Facile handling was possible due to the presence of the PDMS rim structure. **(D)** 3D confocal images of the cells with and without the gel confiner. The reproducibility of the confined space was verified by measuring the cell height. Scale bar = 10 μm. **(E)** Compared with the ‘Not confined’ and ‘Only gel’ systems, the cell height was significantly reduced in the 2D gel confiner cases with smaller variation. n > 90 cells were measured in each case. There was no significant change in the cell height for 24 h under gel confiner conditions. Kruskal-Wallis/Dunn’s multiple comparisons tests were used to compare different sample populations. ***: P < 0.001; N.S.: P > 0.05.

### Slow motility of the DCs triggered by exhaustion

An outline of the DC differentiation process stimulated by LPS for preparation of the xmDCs and the amDCs is shown in **Fig. 2A**. The xmDC and amDC phenotypes were confirmed by flow cytometry and ELISA (**Fig. S1**). The expression levels of the co-stimulatory markers (CD80, CD86, and CD40), the CCR7 chemokine receptor, the antigen-presenting molecules (I-A/I-E), and the DC markers (CD11b, CD11c) were not significantly different for the two phenotypes in terms of the normalised mean fluorescence intensity (MFI) (**Figs. S1A** and **S1B**). To confirm the characteristics of the xmDCs, we performed ELISA to measure the secretion of the pro-inflammatory cytokines IL-10 and IL-12 p70 (**Fig. S1C**). We observed an increased secretion level on Day 1 (24 h), and a decreased secretion level on the following day (48 h) due to DC exhaustion.^17^ In the case of the amDCs, no increased secretion was observed on Days 1 and 2 because of the short LPS stimulation time employed (i.e., <1 h), which is not sufficient to trigger transient cytokine secretion, ^16, 29^ but does alter the up-regulation of antigen-presenting molecules in addition to the cell motility. ^15, 30, 31^

**Figure 2.**
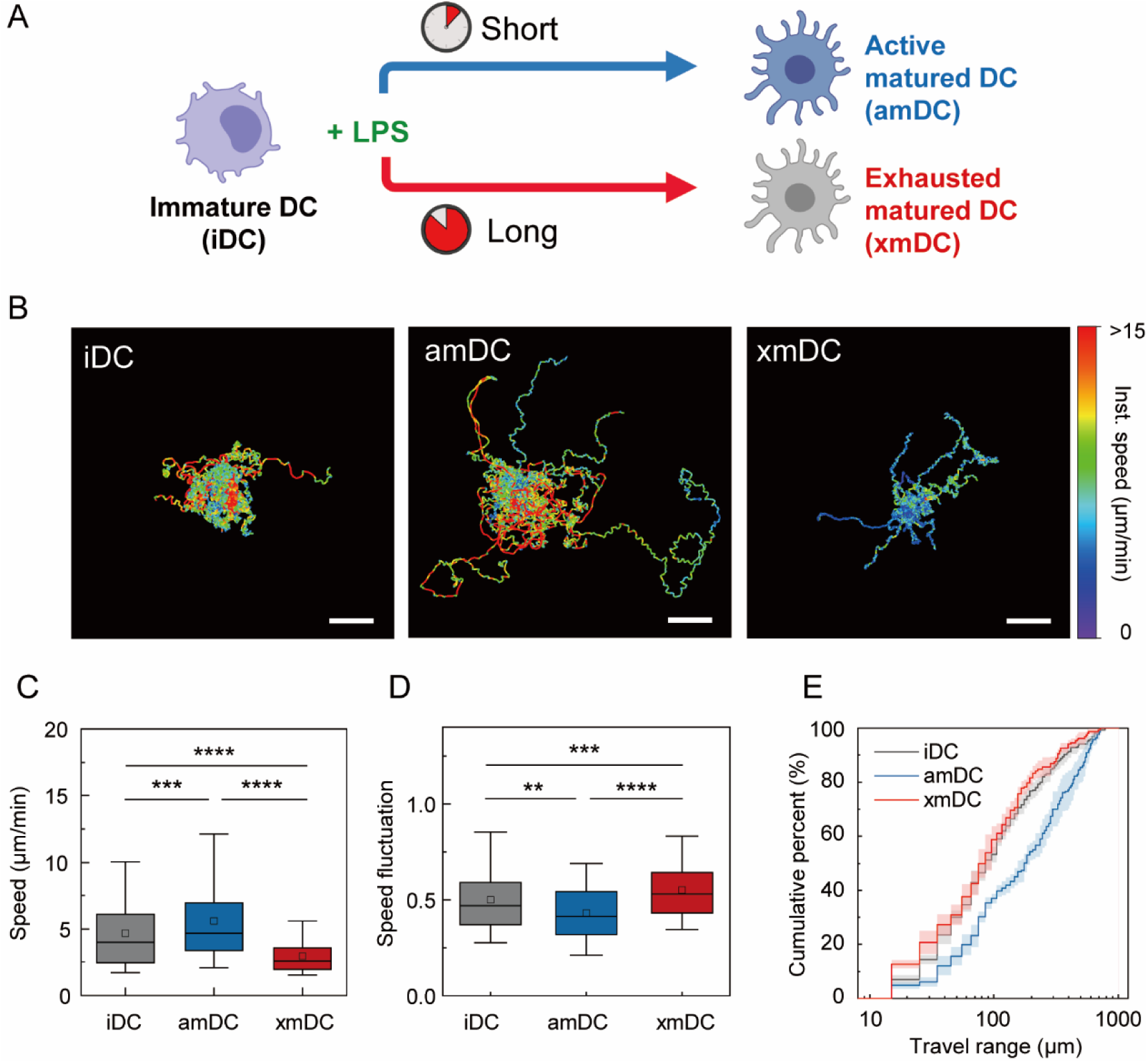
2D Random motility under the agarose gel confiner. **(A)** To differentiate between the amDCs and the xmDCs, different LPS stimulation times were employed. **(B)** Representative trajectories of the iDCs, amDCs, and xmDCs are displayed. The starting point of each trajectory was translated to the origin of the plot. Colours indicate instantaneous speed. One representative experiment out of three is shown. Scale bar = 100 µm. **(C, D)** DC exhaustion induces a slow and more fluctuating motility. **(C)** Mean track speed. **(D)** Speed fluctuation. iDCs: n = 241; amDCs: n = 190; xmDCs: n = 143. Kruskal-Wallis/Dunn’s multiple comparisons tests were used to compare different sample populations; **: p<0.01 ***: p<0.001 ****: p<0.0001. **(E)** Travel range indicating how far DCs can travel. Mean values from 3 independent experiments. The error bar represents the S.D.

To investigate whether exhaustion triggers intrinsic changes in the DC motility, we first identified how these cells move under the agarose gel confinement without the presence of any external factors. As shown in **Fig. 2B** and in Movies **S1–S3**, the xmDCs migrated more slowly than the amDCs and iDCs. More specifically, the xmDCs presented significantly decreased mean track speeds (2.9 ± 0.1 μm/min, mean ±S.E.) compared to the amDCs (5.6 ± 0.2 μm/min) and iDCs (4.7 ± 0.2 μm/min) (**Fig. 2C**). Previously, when iDCs were incubated with LPS, a transiently increased speed fluctuation was observed in the early state.^32^ In the current study, we observed a more significant increased speed fluctuation for the iDCs (0.5 ± 0.0, mean ± S.E.) compared to the amDCs (0.4 ± 0.0). Interestingly, we found that the xmDCs (0.6 ± 0.0) showed a higher speed fluctuation than the iDCs (**Fig. 2D**).

To understand the effect of DC exhaustion on DC travel range, we also measured the distance between track origin to the farthest travel point. In cumulative plot shown in **Fig. 2E**, approximately 60% of xmDC showed the travel range less than 100 μm, similar to iDC, while 40% of amDC travelled to less than 100 μm. In sum, xmDC were slower and more frequently changed their speed than amDC. Thus, DC exhaustion induces distinctive motility pattern change that result in xmDC traveling in a more reduced area compared with amDC counterparts.

To understand the effect of DC exhaustion on the DC travel range, we also measured the distance between the track origin and the farthest travel point. In the cumulative plot shown in **Fig. 2E**, approximately 60% of the xmDCs showed a travel range of <100 μm, which is similar to the case of the iDCs, whereas 40% of the amDCs showed a travel range of <100 μm. These observations indicate that the xmDCs moved more slowly than the amDCs, and changed their speed more frequently. Thus, DC exhaustion induces a distinctive motility pattern change that results in xmDCs travelling over a more reduced area compared with their amDC counterparts.

### xmDC lost persistent motility

MSD is widely used to explain object diffusivity in random migration.^33^ MSD is also used to describe the random motility fof physiological relevance between the cell motility and functional implications, such as the efficiency of space exploration or the searching efficiency.^15, 34-36^ In this study, we found that xmDCs exhibited a reduced persistence compared to amDCs, giving a similar value to iDCs (**Fig. 3A**). It has been previously reported that the diffusivity of amDCs is higher than that of iDCs.^15^ In the current study, we confirmed similar trends. In addition, we found that the diffusion coefficient and exponent α of the xmDCs were significantly smaller than those of the amDCs, but similar to those of the iDCs (**Figs. 3B** and **3C**).

**Figure 3.**
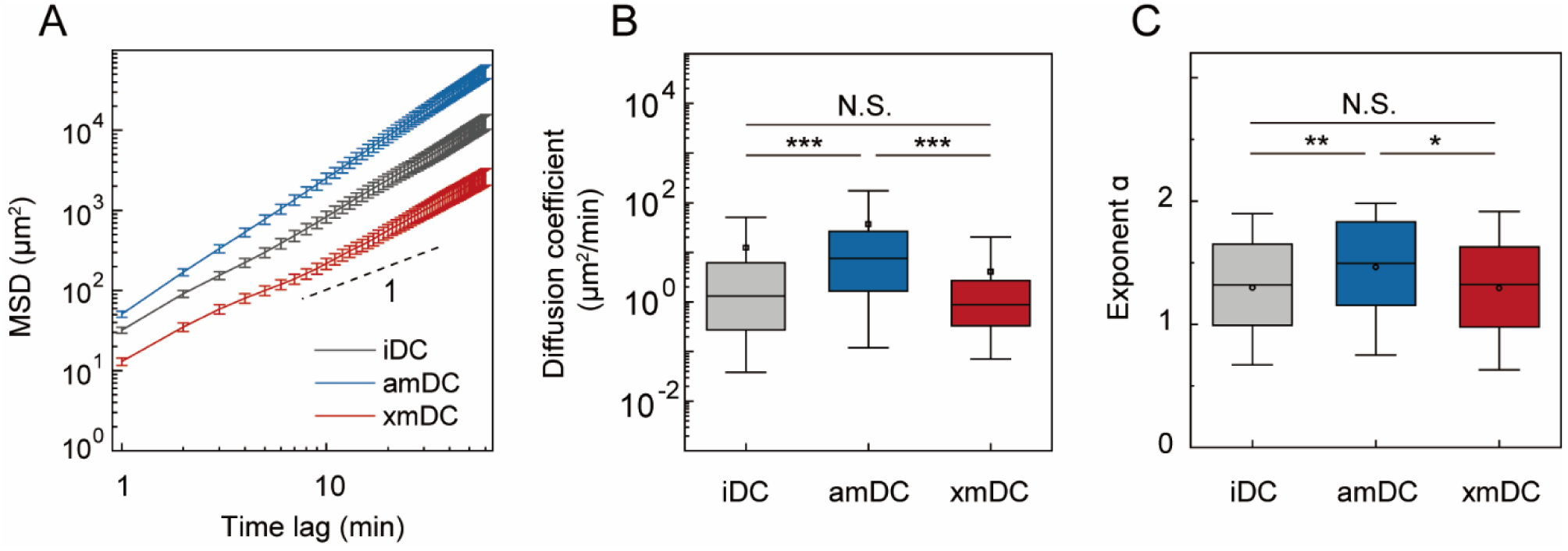
Decreased diffusivity of xmDC. **(A)** The mean squared displacement (MSD) was plotted to demonstrate the persistent random motility of the DCs. Lines indicate the mean values of the MSD from three independent experiments, whereas the error bar represents the standard deviation (S.D.). **(B, C)** MSD curves of the various trajectories were collected from τ = 6–30 min and were fitted to the power law form to calculate **(B)** the diffusion coefficient and **(C)** exponent α. Data were pooled from 3 independent biological replicates. iDC: n = 167; amDC: n = 148; xmDC: n = 99. Kruskal-Wallis/Dunn’s multiple comparisons tests were used to compare different sample populations; N.S.: p > 0.05; *: p < 0.05; **: p < 0.01; ***: p < 0.001.

Subsequently, we analysed the turning angle distribution. More specifically, the turning angle was measured at each time lag in the path (**Fig. 4A**) and was plotted on a heat map (**Fig. 4B**), which indicates the change in direction of the DCs with different time lags. It was found that for the amDCs, the fraction of turning angles <30° was evenly distributed between 2 and 12 min, whereas for the xmDCs, a varied distribution was observed. More specifically, the xmDCs presented an increased fraction between 150° and 180° at 3 min compared to the amDCs, but this difference disappeared at 12 min (**Fig. 4C**). In addition, the turning angle distribution of the xmDCs was found to be similar to that of the iDCs (**Fig. S2**), thereby indicating that the xmDCs change direction after approximately 3 min, as in the case of the iDCs. In contrast, the amDC turning angle distribution was rather persistent. These results consistently support the suggestion that xmDCs lose their persistent motility.

**Figure 4.**
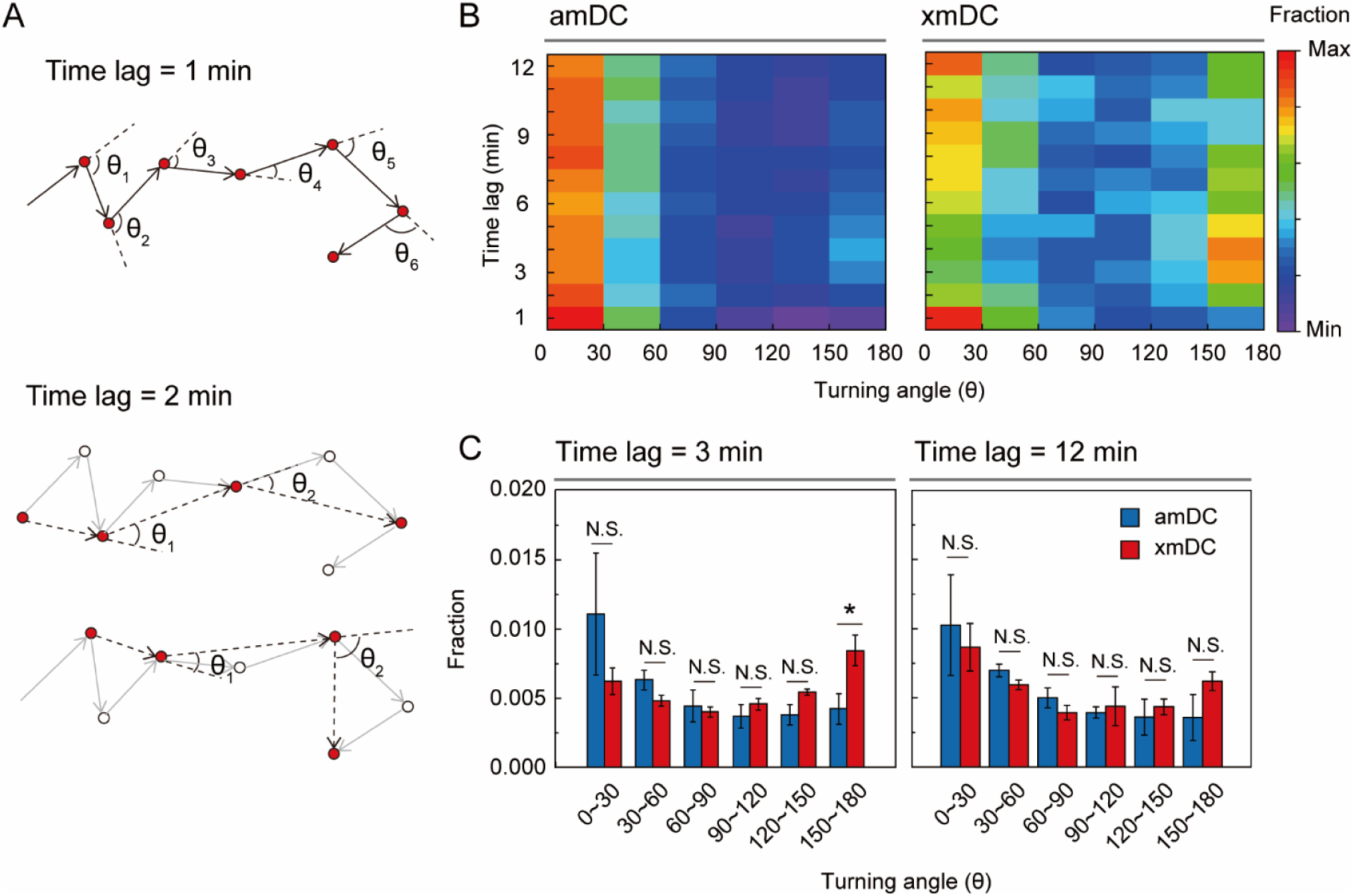
The turning angle distribution of the xmDCs shows non-persistent motion. **(A)** Scheme outlining measurement of the turning angles with different time lags. **(B)** Turning angle distribution based on the time lag. Each rectangle depicts the fraction of the turning angle (bin size = 30°), and is indicated according to its relevant colour code. **(C)** Detailed turning angle distributions at time lags of 3 and 12 min. The un-paired two-tailed Student’s t-test was used to compare different sample populations. N.S.: P < 0.05 and *: P > 0.05.

### Preservation of CCL19 chemotaxis in xmDCs

Chemotaxis against the CCL19 concentration gradient is an important phenomenon for initiating adaptive immunity and recruiting immune cells ^37^ As mentioned above, we found that the intrinsic motility of xmDCs becomes slow and less persistent when there is no directional cue. Based on these observations, we examined whether exhausted motility also appears under CCL19 chemotaxis. More specifically, we designed a confined microchannel with a concentration gradient to study the chemotaxis of xmDCs. To achieve a stable concentration gradient, CCL19-embedded collagen was employed to support the diffusion control taking place at the endpoint of the confined microchannel (**Fig. 5A**). The stability of the concentration gradient was determined by the diffusion of fluorescence-conjugated dextran (CF488A-dextran, 10 kDa) over time (**Fig. 5B**), wherein a stable gradient was maintained between 5 and 15 h in the 1D microchannels (width: 20 μm, length: 1000 μm, height: 3 μm).

**Figure 5.**
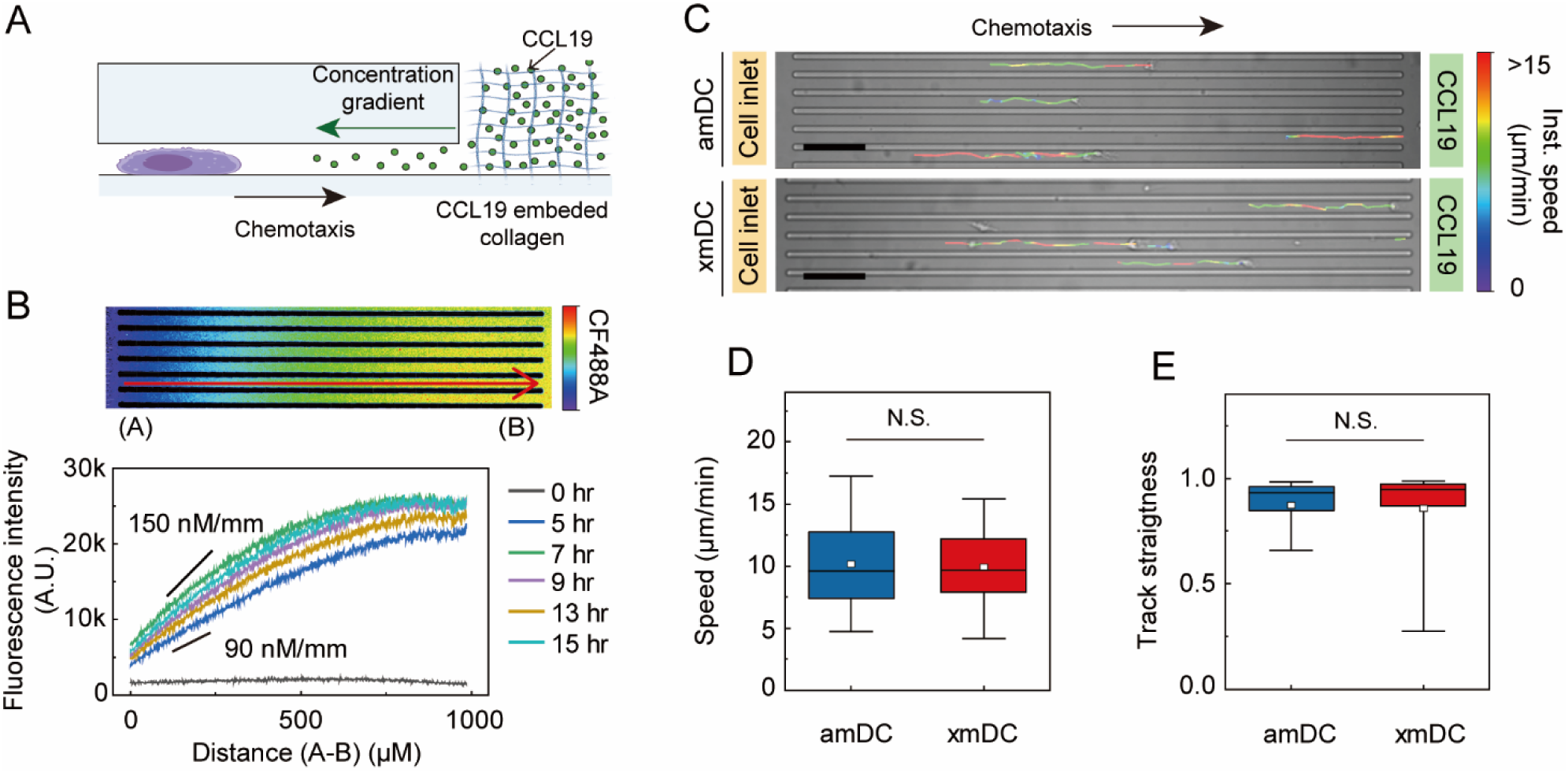
Chemotaxis of xmDCs under a CCL19 concentration gradient. **(A)** CCL19-embedded collagen located at the endpoint of the confinement channel. CCL19 diffused from collagen to the confinement channel. **(B)** The stability of the concentration gradient was confirmed by testing with CF488A-dextran (10 kDa). A representative image of the concentration gradient is displayed in the upper panel. The fluorescence intensity profile over time based on the upper panel is presented in the lower panel. One representative result out of five is presented. The gradient was maintained from 5–15 h. The gradient scale was maintained up to approximately 150 nM/mm and the initial gradient scale was >90 nM/mm. **(C)** Representative image of DC migration under a CCL19 concentration gradient. The colour code depicts the instantaneous speed of the migrating DCs. Scale bar = 100 µm. **(D)** Mean track speed, and **(E)** the track straightness of chemotaxis. The Mann-Whitney test was used to compare different sample populations. N.S.: p > 0.05. (amDC, n = 165; xmDC, n = 138). All figures are based on pooled data from 3 independent biological replicates.

Interestingly, both the xmDCs and the amDCs exhibited directional motility in a confined microchannel (**Fig. 5C**), and no significant differences were observed between the mean track speeds of the two phenotypes (i.e., xmDCs: 9.9 ± 0.3 μm/min, mean ± S.E.; amDCs: 10.2 ± 0.3 μm/min, **Fig. 5D**). Similar values were also observed in terms of the track straightness (i.e., xmDCs: 0.9 ± 0.0, amDCs: 0.9 ± 0.0, mean ± SE, **Fig. 5E**). In the absence of CCL19, neither the xmDCs nor the amDCs exhibited chemotaxis. Considering these results, DC exhaustion did not have a significant impact on CCL19 chemotaxis, and the fact that there was no significant difference in the expression of CCR7 also supports this result (**Fig. S1A**).

## Discussion

DC motility is essential to ensuring an appropriate immune response, both in terms of searching for antigens through innate immunity and initiating adaptive immunity. In this regard, DC exhaustion is an essential topic to understand chronic infection or inflammation, but its impact on cellular locomotion had not been extensively studied prior to the current study. To address this gap in the literature, we quantified the motility of individual cells under agarose gel confinement to determine the effect of exhaustion on DC motility. We found that xmDCs moved slowly with fluctuating speeds, and their travel range became narrower. In addition, xmDCs presented a lower diffusion coefficient and a smaller exponent α than amDCs, suggesting that xmDCs exhibit a slower and less persistent motility than that observed for amDCs. This observation was also confirmed by the distracted turning angle distribution observed for the former. These results therefore indicate that DC exhaustion can result in both a reduction in cytokine secretion and a lower intrinsic cellular motility.

In a previous study, the incubation of DCs with LPS (100 ng/mL) for 0.5 h was found to be sufficient to upregulate CD86 co-stimulatory molecules and CCR7 expression, which resulted in DC activation.^15^ These activated DCs obtained using a short LPS pulse exhibited fast and persistent intrinsic migration in the absence of chemokines, which could be interpreted as a strategy to reach the lymphoid organs more efficiently.^15^ In contrast, the relative mean speed of iDCs incubated with LPS (10 μg/mL) for ≥2 h was found to reduce their motility,^32^ thereby implying that the presence of LPS inhibits the active motility of DCs.^32^ Interestingly, our results showed that xmDCs upregulate antigen-presenting molecules to a similar extent as amDCs, but their intrinsic motility remains slow, even when the LPS is removed. Based on the observation of these distinctive motility phenotypes, it could be assumed that DC exhaustion leads to a unique motility pattern of slow, fluctuating, and narrower spatial spanning.

The persistent chemotaxis towards the CCL19 concentration gradient is a characteristic of mDC motility. However, it is not known whether the exhaustion of DCs influences their chemotactic motility. Although we found that their intrinsic motility pattern was altered upon DC exhaustion, the chemotactic motility did not differ between xmDCs and amDCs. These results could be expected from the similar expression levels of CCR7 between the two cases, thereby allowing us to conclude that DC exhaustion alters the random motility but does not have a significant impact on CCL19 chemotaxis.

Due to the fact that the in vivo microenvironment of inflammatory tissues or chronic infection sites is extremely complex and dynamic, single-cell tracking technology using microfluidic devices with well-controlled parameters that reproduce the tissue microenvironment is an important tool to expand our knowledge of intercellular communication and its role in cellular motility and cellular functions. We therefore believe that further studies are required to gain a better understanding of the influence of dynamic changes in both physical and chemical stimuli to mimic chronic infection sites in tissues. For example, inflammatory tissue presents a diverse range of tissue stiffness degrees depending on the disease state. ^38^ Moreover, the DC motility under topological differences such as extracellular matrix alignment could also be considered to reproduce tissue-specific models.^39^ It has been demonstrated that several inflammatory cytokine control the fast motility of DCs.^40, 41^ Considering the unique cytokine interactions taking place during maturation, the adaptive DC motility attributed to inflammatory cytokines should be further studied.

## Conclusions

In this study, we investigated the effect of dendritic cell (DC) exhaustion on cellular motility using microfluidic devices that allowed precise control of the cell confinement. More specifically, we studied the cellular migration of exhausted DCs in tissue-mimicked confined environments. It was found that the speed of exhausted matured DCs (xmDCs) decreased significantly compared to amDCs and iDCs. In contrast, the speed fluctuation increased compared to that of amDCs and was similar to that of iDCs. In terms of the diffusivity, it was found that the diffusivity of the xmDCs was significantly lower than that of the amDCs, thereby implying that DC exhaustion reduces the space exploration ability. Importantly, the obtained results indicate that CCR7-dependent chemotaxis against CCL19 in xmDCs was not considerably different from that observed in amDCs. Overall, a unique intrinsic cell migration behaviour was observed for xmDCs, which results in the DCs remaining at the site of infection, although a well-preserved CCR7-dependent chemotactic motility was also maintained. Finally, our results provide additional insight into how DCs respond and control their motility in chronic infection and related disease, which is of particular importance because DC motility in the confined environment of peripheral tissues is key to ensuring an efficient immune sentinel function.

## Supporting information

Electronic Supplementary Information

## Author Contributions

Y.C. and Y.K.C. designed the research; Y.C. and V.S. performed the experiments. Y.C. and Y.L. performed cell motility tracking. Y.C. analysed and interpreted all the data, and Y.C., V.S., and Y.K.C. wrote the paper. All authors approved the final version of the manuscript.

## Acknowledgements

This work was supported by the Institute for Basic Science (IBS-R020-D1) funded by the Korean Government. We thank Ms. Jaeun Kwon for supporting the tracking. We appreciate the technical support of Mr. Jin-Hoe Hur, UNIST-Optical Biomed Imaging Center (UOBC) for helping with confocal imaging.

