## Supplementary material for "Exhausted mature dendritic cells exhibit a slower and less persistent random motility but retain chemotaxis against CCL19": Electronic Supplementary Information

**The Supplementary Information file includes:**

- Supplementary Materials and Methods
- Supplementary Figures S1 – S2
- Supplementary Movies S1 – S5

### Supplementary Materials and Methods

#### Preparation of bone-marrow derived DC (BMDC)

The tibias and femurs from BALB/c mice (8–12 weeks, female) were flushed and red blood cells were lysed with ammonium-chloride-potassium (ACK) lysis buffer (Gibco, Thermo Fisher Scientific). Bone marrow cells were plated on 24 well tissue culture plates (1 × $10^6$ cells/mL) in complete medium (RPMI 1640, supplemented with 5% (v/v) fetal bovine serum (FBS), 1% (v/v) antibiotic–antimycotic solution, 1% (v/v) HEPES buffer, 0.1% (v/v) 2-mercaptoethanol, and 20 ng/mL recombinant mouse granulocyte macrophage colony stimulating factor (GM-CSF) (Peprotech). The medium was replaced with fresh complete medium containing GM-CSF every 2 days. On day 6, non-adherent and loosely adherent cells were collected by gentle pipetting and transferred to Petri dishes.

#### Antibodies used in flow cytometry analysis

The following antibodies were used for the flow cytometry experiments: CD86-FITC (clone GL1, 1:200), CD40-PE (clone 1C10, 1:200), MHCII I-A/I-E-FITC (clone M5/114.15.2, 1:200), and CCR7-PE (clone 4B12, 1:200) from Thermo Fisher Scientific, and CD11b-APC (clone M1/70, 1:200), CD11c-APC (clone HL3, 1:200), and CD80-PE (clone 16-10A1, 1:200) from BD Biosciences. The acquired data were analysed using FlowJo (BD).

#### Cytokine measurements

Conditioned media for cytokine quantification were prepared as described below. For the xmDCs, the iDCs (3 × $10^5$ cells/mL) were incubated with 100 ng/mL LPS. For the amDCs, the iDCs were stimulated with LPS for 0.5 h. The LPS-containing medium was carefully washed three times with fresh complete medium and incubated with the same number of xmDCs (3 × $10^5$ cells/mL). The iDCs were incubated with fresh complete medium. The cell-conditioned medium was collected after 24 h, and the harvested cells were cultured again with fresh conditioned medium for a following 24 h (day 2). The concentrations of cytokines IL-10 and IL-12 p70 in the conditioned media were determined using DY417 and DY419 ELISA DuoSets (R&D Systems), respectively.

#### Fabrication of agarose gel confiner

Mixtures of PDMS (Sylgard 184 Silicone Elastomer Kit, Dow Corning) with a 10:1 ratio (weight ratio, w/w prepolymer to curing agent) was used to prepare main body and a 30:1 ratio was for sticky PDMS bottom. Prior to casting the gel solution, the PDMS structure was placed on a Petri dish. Subsequently, 2.4% (w/v) low-melting agarose was dissolved in phenol red-free HBSS buffer (Gibco) at 90 °C for 20 min. The melted agarose solution was slowly cooled to 40 °C, and the same volume of warm (37 °C) 2× conditioned medium (RPMI 1640, 10% (v/v) FBS, 2% (v/v) HEPES buffer, 2% (v/v) antibiotic–antimycotic solution, 0.2% (v/v) 2-mercaptoethanol) was mixed thoroughly to give a final agarose gel concentration of 1.2% (w/v), which was cast onto the PDMS structure. Curing was then carried out for 20 min at room temperature, and the cured structure was stored in complete medium and incubated overnight in a cell culture incubator.

#### Measurement of the cell height

The heights of the fluorescence-labelled DCs under confinement were measured to determine the reproducibility of the confined environment using laser scanning confocal microscopy (A1R, Nikon). The DC suspension was stained with DiO (Invitrogen) seeded on the 20 μg/mL FN-coated glass substrate and incubated for 20 min before being covered by a gel confiner. The assembled structure was placed on a Chamlide magnetic chamber (Live Cell Instruments) for imaging. In the 'Not confined' case, the DCs were seeded on a glass substrate following the same process, and an aliquot (1 mL) of complete medium was carefully added. For the ‘Only gel’ case, the agarose gel solution (1 mL) was cast and cured in a Chamlide magnetic chamber (Live Cell Instruments) and incubated with complete medium overnight. Subsequently, the DCs seeded on the substrate were carefully covered with cured agarose gel in a Chamlide magnetic chamber after the removal of any remaining complete medium. The 3D confocal images were acquired using a 100× Plan Apo lens, with a z interval of 0.5 μm. More than 20 cells were measured in five different positions from nine independent samples. The cell height was manually measured using NIS Elements (Nikon).

**DC motility analysis**

In the agarose gel confinement experiment, brightfield maximum intensity projection (MIP) was performed to track the non-labelled sample. More specifically, the sample focal plane was focused and two more image sequences were taken along the z-axis. Subsequently, rolling ball background subtraction was performed on the DIC image, and the Z-stacks were overlaid using NIS-Elements (Nikon). The cell trajectories were automatically detected using the Imaris (Bitplane) 'Spots' function. Cells were determined using intensity thresholding and the particle size was estimated to be 20 µm in diameter. The position of the centre of mass was tracked, and the tracking errors were corrected manually. Previous studies applied reasonable filtering to the raw data to remove artifacts, noise, and dead or dying cells when dealing with trajectory analysis for DC migration.[1] Similarly, we also used three filtering criteria in this work, which are related to the mean track speed (≥1.5 µm/min), the track duration (≥60 min), and the maximal distance from its starting position (≥20 µm).

**Preparation of chemotaxis chip**

For device production, PDMS composed of a 10:1 ratio of elastomer:curing agent was poured over the micro-patterned moulds, degassed, and cured at 65 °C overnight. Subsequently, the structures were carefully peeled off and cut. Holes measuring 3 mm in diameter were punched on opposite sides of the structure to accommodate the CCL19-embedded collagen and the DC suspension. The trimmed PDMS structure was then plasma-activated and bonded to glass slides at 65 °C for 10 min. Subsequently, the structure was sterilised by UV irradiation for 10 min, and the inner space was coated with 20 µg/mL FN for 1 h and then washed three times with PBS before loading the CCL19-embedded collagen. The opposite hole was loaded with 200 nM CCL19 (PeproTech) embedded in 2.4 mg/mL type-I bovine collagen (Purecol®, Advanced Biomatrix) to achieve a concentration gradient. Subsequently, the structure was incubated at 37 °C for 30 min and then washed three times with complete medium.

### Supplementary Figures

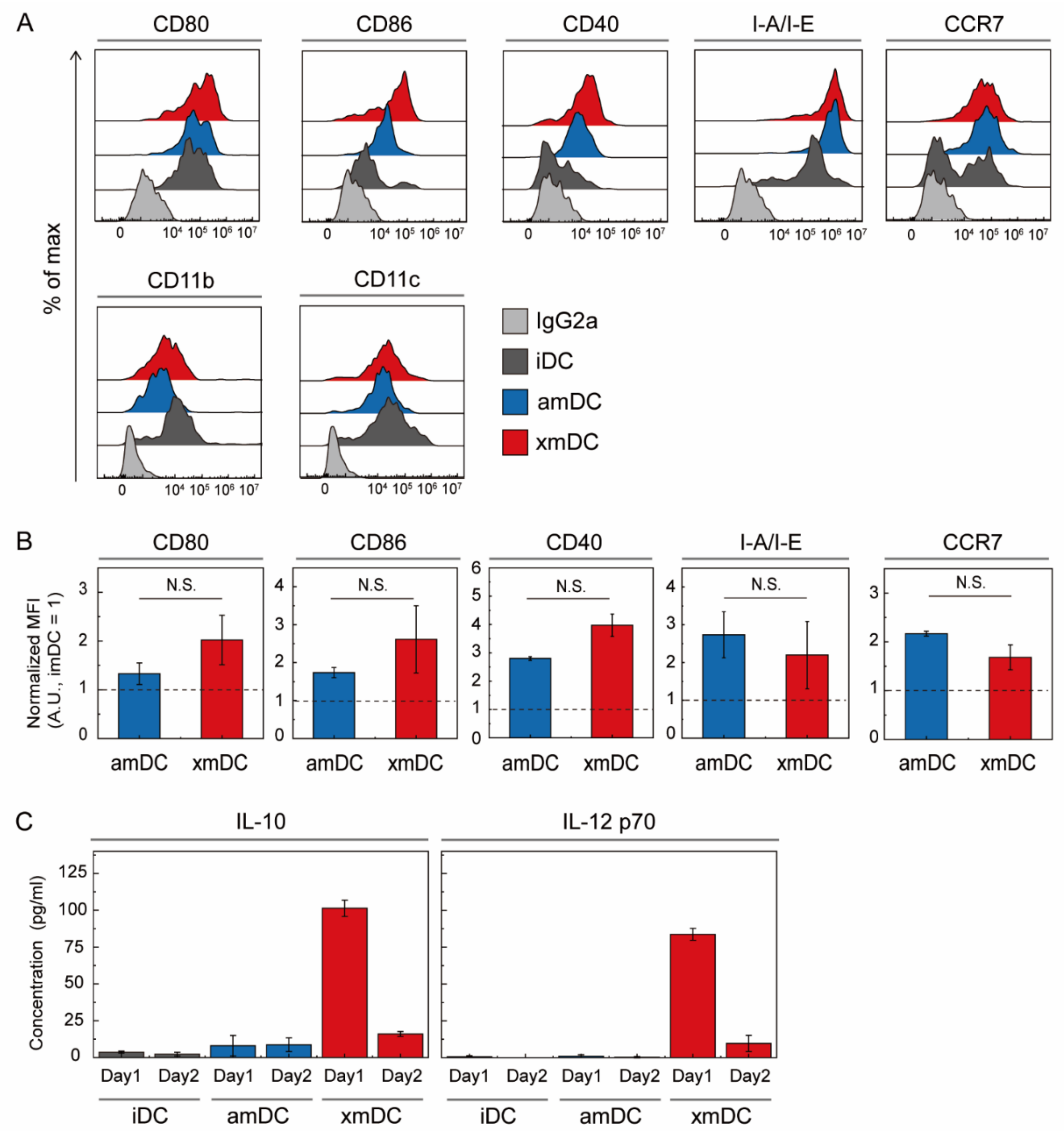


**Figure S1. Characteristics of the xmDCs.** To differentiate between exhausted and mature phenotypes, iDCs were stimulated by LPS (100 ng/mL) for 24 h (xmDC) or 0.5 h prior to incubation with fresh medium for 24 h (amDC). (**A**) Representative flow cytometry data of DC maturation and the DC marker. One representative experiment out of three is shown. (**B**) Normalized mean fluorescence intensity (MFI) of the DC maturation marker. Each value was normalized based on the mean of the iDCs. No significant difference exists between the amDCs and the xmDCs. The un-paired two-tailed Student's t-test was used to compare different sample populations. N.S.: $P > 0.05$, $N = 3$. (**C**) Cytokine secretion levels of IL-10 and IL-12 p70. Cell culture supernatants from the amDCs and the xmDCs were collected at Day 1 (0–24 h). After 24 h, the culture media were exchanged with fresh medium at the beginning of Day 2 (24–48 h) and harvested. The amounts of IL-10 and IL-12p70 secreted by the DCs were measured by the enzyme-linked immunosorbent assay (ELISA) approach. Comparison of the data for Days 1 and 2 suggest a loss of the cytokine secretion capacity.

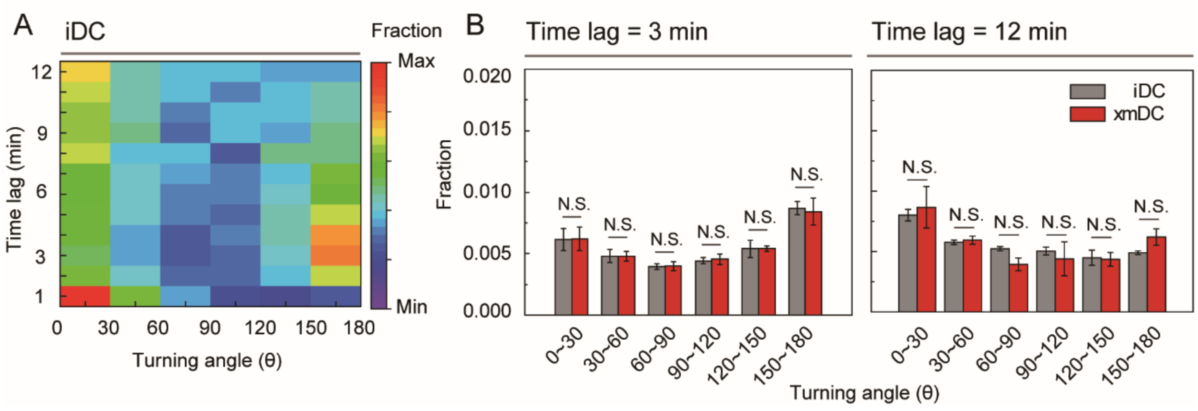


**Figure S2. Turning angle distributions of the xmDCs and the iDCs**. (**A**) Variation in the turning angle distribution with the time lag. Each rectangle depicts the fraction of the turning angle (bin size = 30°), which is indicated by the relevant colour code. (**B**) Detailed turning angle distributions at time lags of 3 and 12 min. The un-paired two-tailed Student's t-test was used to compare different sample populations. N.S.: P > 0.05, N = 3.

### Supplementary Movies

**Movie S1:** iDC Motility under agarose gel confiner and time-lapse during brightfield microscopy (10×) at a frequency of 1 image/min. Scale bar = 50 µm.

**Movie S2:** amDC Motility under agarose gel confiner and time-lapse during brightfield microscopy (10×) at a frequency of 1 image/min. Scale bar = 50 µm.

**Movie S3:** xmDCs Motility under agarose gel confiner and time-lapse during brightfield microscopy (10×) at a frequency of 1 image/min. Scale bar = 50 µm.

**Movie S4:** amDC Migration along a CCL19 gradient in 20 µm (w) × 3 µm (h) fibronectin-coated micro-channels and time-lapse during brightfield microscopy (10×) at a frequency of 1 image/min. The source of CCL19 is at the right side of the movie. The colour indicates the instantaneous speed of the track, and the speed ranges from 0 to >15 µm/min. Scale bar = 100 µm.

**Movie S5:** xmDC Migration along a CCL19 gradient in 20 µm (w) × 3 µm (h) fibronectin-coated microchannels and time-lapse during brightfield microscopy (10×) at a frequency of 1 image/min. The source of CCL19 is at the right side of the movie. The colour indicates the instantaneous speed of the track, and the speed ranges from 0 to >15 µm/min. Scale bar = 100 µm.
